# Identification of a lineage-agnostic splicing signature caused by PRMT5 inhibition

**DOI:** 10.64898/2026.03.26.714409

**Authors:** Matthew R. Tonini, Samuel R. Meier, Shangtao Liu, Kevin M Cottrell, John P Maxwell, Jannik N. Andersen, Alan Huang, Luisa Cimmino, Kimberly J Briggs

## Abstract

PRMT5 is a type II arginine methyltransferase that forms an active complex with methylosome protein WDR77 (MEP50) to catalyze the symmetric dimethylation (SDMA) of arginine residues in proteins that regulate biological roles including apoptosis, DNA damage response and RNA processing. Some of the best characterized PRMT5 substrates are the small nuclear ribonucleoproteins SNRPB, SNRPD1 and SNRPD3, which are critical for spliceosome assembly and RNA splicing fidelity. MTAP-deleted cancers exhibit increased sensitivity to PRMT5 inhibition due to elevated levels of methylthioadenosine (MTA), a natural inhibitor of PRMT5. This vulnerability is exploited by MTA-cooperative PRMT5 inhibitors, exemplified by TNG908 and TNG462 which selectively target PRMT5 in MTAP-deleted cells while sparing MTAP-wildtype (WT) cells. Consistent with this mechanism, treatment with TNG908 in preclinical studies induces widespread splicing alterations in MTAP-deleted cancer models, with minimal effects in MTAP-WT cells. These splicing changes are consistent across diverse MTAP-deleted tumor types, including glioblastoma, pancreatic, and non-small cell lung cancer, indicating a histology-agnostic response to PRMT5 inhibition. Moreover, treatment of MTAP-WT cells with exogenous MTA mimics the splicing alterations observed with PRMT5 inhibition, as does pharmacologic inhibition of MTAP further supporting a mechanistic link between MTA accumulation, PRMT5 modulation, and aberrant splicing. Given that MTAP deletions occur in approximately 10–15% of human cancers, the identification of a robust RNA splicing signature offers a valuable pharmacodynamic biomarker for monitoring the activity of PRMT5 inhibitors. This splicing-based readout may also serve as a predictive biomarker of therapeutic response, offering greater specificity than global SDMA levels. Collectively these data suggest that a PRMT5-dependent RNA splicing signature can monitor the pharmacodynamic activity of MTA-cooperative PRMT5 inhibitors in MTAP-deleted cells.

## INTRODUCTION

The enzyme methylthioadenosine phosphorylase (*MTAP*) holds a unique position as the singular characterized enzyme capable of metabolizing 5’-methylthioadenosine (MTA) within the methionine and adenine salvage pathways. Frequently, *MTAP* is co-deleted with the tumor suppressor *CDKN2A* due to their proximity on chromosome 9p21 (1, 2). Consequently, metabolomic and biochemical experiments established that the absence of MTAP leads to an accumulation of intracellular levels of MTA, which inhibits the methyltransferase activity of type II protein arginine methyltransferase 5 (PRMT5) through a S-adenosyl methionine (SAM) competitive manner (3). PRMT5 forms a catalytically active complex with WDR77 (MEP50) and catalyzes symmetric dimethylation (SDMA) of arginine residues in a wide array of substrates (4, 5). These include histones (H2A, H3, H4) and non-histone targets such as p53, E2F1, NM23, c-Myc, and ST7, thereby regulating chromatin structure and gene expression (4, 5). Beyond its role in epigenetic regulation, PRMT5 participates in the methylation of the RNA-binding proteins SmB/B′, SmD1, and SmD3 affecting their function in RNA processing, splicing, and transport (4). This highlights PRMT5’s critical role in maintaining transcriptomic integrity.

Multiple groups have identified an intrinsic vulnerability in *MTAP*-deleted cells, where additional perturbation of PRMT5 markedly reduces the viability of *MTAP*-deleted cancer cells (1–3). This genetically defined sensitivity is a classic example of collateral lethality, where the loss of a non-essential passenger gene (*MTAP*) creates a dependency on a functionally related target (PRMT5) (6). As a result, focus has shifted toward developing biomarkers to support clinical translation. *MTAP* deletion is now commonly used as a patient selection biomarker in trials of MTA-cooperative PRMT5 inhibitors, while SDMA loss, a direct consequence of PRMT5 inhibition, serves as a validated pharmacodynamic biomarker for PRMT5 target engagement (7–10).

While SDMA loss is widely used as a pharmacodynamic biomarker of PRMT5 inhibition, it presents several limitations. As a global marker of arginine methylation, SDMA does not capture the downstream functional consequences of PRMT5 inhibition on specific cellular processes. Moreover, SDMA levels may not sensitively distinguish between partial and complete inhibition or reflect the context-dependent effects seen in *MTAP*-deleted versus *MTAP*-WT tumors. These constraints are particularly relevant given the emergence of both MTA-cooperative and non-MTA-cooperative PRMT5 inhibitors in clinical development, which may differ in their on-target dynamics and tumor selectivity. Consequently, there is an unmet need for alternative pharmacodynamic biomarkers that more directly reflect the cellular impact of PRMT5 inhibition and may improve patient stratification and therapeutic monitoring.

Mechanistically, *MTAP* loss results in MTA accumulation, which partially inhibits PRMT5 and lowers the threshold for pharmacological inhibition. Treatment of *MTAP*-WT cells with exogenous MTA recapitulates PRMT5-dependent phenotypes, reinforcing MTA’s role as a key mediator of synthetic lethality in *MTAP*-deleted cells (1–3). Accordingly, the therapeutic activity of MTA-cooperative PRMT5 inhibitors depends on sufficient intratumoral MTA accumulation, which creates a distinct metabolic context in *MTAP*-deleted tumors. Characterizing this metabolically driven vulnerability could enable identification of optimal patient subsets and enhance biomarker-driven clinical trial design.

Among the functional consequences of PRMT5 inhibition, disruption of RNA splicing has emerged as a prominent and measurable effect. Alternative splicing (AS) allows a single gene to produce multiple mRNA isoforms through selective exon inclusion or exclusion, greatly expanding proteome diversity and enabling fine-tuned gene regulation (11–15). Aberrant splicing has been considered a promising new avenue for biomarker development due to the significant correlation between splicing and various tumor types (9, 10). Because splicing governs the expression and function of many essential genes, changes in splicing patterns may also serve as a sensitive readout of therapeutic response.

An expanding body of research underscores that inhibiting PRMT5 triggers thousands of alternative splicing events, with a notable enrichment in genes associated with RNA processing, DNA damage response, and cell cycle regulation. When *MTAP* is deleted, MTA can accumulate to levels sufficient to partially inhibit PRMT5, which may lead to defective spliceosome activity. Therefore, we surmised that an RNA splicing-based signature could function as a quantitative surrogate for MTA accumulation and a functional biomarker of PRMT5 inhibitor activity. Using RNA sequencing we analyzed the effects of PRMT5 inhibition on alternative splicing in various tumor lineages. We identified and characterized a subset of these alternative events and found them to be lineage agnostic and PRMT5 dependent. Furthermore, treatment of *MTAP*-WT cells with exogenous MTA partially perturbed PRMT5 activity leading to increased splicing alterations, consistent with the known mechanism of PRMT5 inhibition. We also demonstrate that modulation of this splicing signature can be leveraged to monitor PRMT5 inhibitor efficacy in both in vitro and in vivo models. We propose that this splicing signature represents a quantitative and mechanistically supported pharmacodynamic biomarker of PRMT5 inhibitor activity. In contrast to static methylation-based markers such as SDMA, splicing-based biomarkers may offer enhanced sensitivity and biological relevance, particularly in the context of partial PRMT5 inhibition by MTA-cooperative compounds. As such, incorporating splicing signatures into clinical development may improve real-time monitoring of drug activity, guide dose optimization, and support mechanism-based patient stratification in trials of both MTA-cooperative and non-cooperative PRMT5 inhibitors.

## METHODS

### Reagents

TNG908 and AG-270 (custom synthesis) (16, 47). 5’-S-Methyl-5’-thioadenosine (Sigma Aldrich #D5011), MTDIA (#HY-101496), Gemcitabine (#S1149), Docetaxel (#S1148), and Palbociclib (#S1116) inhibitors purchased from Selleck Chem and MedChemExpress where available. Anti-glyceraldehyde-3-phosphate dehydrogenase (GAPDH), Anti-Tubulin, Anti-SDMA, Anti-PRMT5, and Anti-MTAP antibodies were from Cell Signaling Technology (Danvers, MA).

### Cell culture

Human cancer cell lines: skin cutaneous melanoma (SK-MEL-5, A101D, Hs294T, IGR-1, SH-4), glioblastoma (LN18, GB-1, U-87 MG, AM-38, KS-1, A172), pancreatic adenocarcinoma (Panc03.27, Panc10.05, SU.86.86, BxPC-3, Miapaca2), mesothelioma (ONE58, SDM103T2, NCI-H2052, ZL5, ACC-MESO-4), cholangiocarcinoma (KKU-100), lung adenocarcinoma (A549, NCI-H1373, NCI-H520, SK-LU-1, HARA, NCI-H322, HCC-1588, SW-900, HCC4006, NCI-H1650, LU99, SW1573), colorectal adenocarcinoma (HCT116) and chronic myeloid leukemia (HAP-1) cells were obtained from ATCC (Manassas, VA) and cultured in cell dependent media (Invitrogen, Carlsbad, CA) supplemented with 10% fetal bovine serum (Hyclone, Logan, UT).

### Western Blot

Cells were washed with cold PBS and then lysed in RIPA buffer (Cell Signaling Technology). After centrifugation at 14,000 rpm for 15 min at 4 °C, the supernatants were collected, and the protein concentrations were measured using bicinchoninic acid protein assay reagent (Bio-Rad). Subsequently, equal amounts of proteins were separated in NuPAGE 4 to 12% Bis-Tris gradient gel (Invitrogen NP0335), and transferred onto nitrocellulose membranes (Invitrogen B301002). After blocking with 5% milk, the membranes were then probed at 4 °C overnight with various primary antibodies, the next morning membranes were washed with TBST (20 mM Tris, 150 mM NaCl, 0.1% Tween 20; pH 7.6), and incubated with horseradish peroxidase (HRP)-conjugated secondary antibodies (Promega) at room temperature for 1 h. Finally, after washing with TBST, the antibody-bound membranes were treated with enhanced chemiluminescent Western blot detection reagents (GE Healthcare) and visualized with an X-ray film (GE Healthcare).

### Apoptosis

Cells were seeded into 96-well plates (Sigma #CLS4594) (3000 cells/well) and treated with specified compounds at specified concentrations for 3-days. Caspase-Glo 3/7 Assay System (Promega #G8091) was used for measuring Caspase 3/7 under different conditions according to the manufacturer’s protocol. Measurement was determined by luminescence using an EnVision XCite 2105 Multimode Plate Reader (PerkinElmer).

### Splicing Assay

RNA was extracted from cell pellets using RNeasy Plus Mini Kit (50) from Qiagen (#74134) as per manufacturer instructions. Eluted RNA was quantified by Nanodrop 8000 (ND8000P21H2). Flanking exon RT-PCR, in which the primers anneal to the exons upstream and downstream of the exon of interest was performed using QIAGEN OneStep RT-PCR Kit (#210212). RT-PCR was performed as an endpoint on 100ng of extracted RNA with the following cycling conditions to amplify the alternatively spliced transcripts: 50C for 30min, 95C for 15 min, 40 cycles of 94C for 60s, 58C for 30s and 72C for 1 min, followed by 72C for 10 min. Expression levels of each exon were resolved on 2% agarose gels visualized with SYBR Safe DNA Gel Stain from ThermoFisher (#S33102) and quantified using Bio-Rad imaging software. ACTB levels were used as an internal reference for RT-qPCR analysis. All the primers and expected amplicon sizes are listed in the Supplementary Table.

### RNA-Seq and Library Preparation

LN18 MTAP-deleted cells were treated with either DMSO or TNG-0239908 at cell line specific IC_50_ (0.2µM) for 72 hours. Total RNA was purified using RNeasy Plus Mini Kit (Qiagen #74136) following manufacturer’s instructions. RNA purity was determined to be between 1.8-2.2A based on A260/280 readings. Sample mRNA was isolated by Poly(A) mRNA magnetic isolation and RNA-Seq libraries were prepared and assessed for quality at Azenta Life Sciences. Sample sequence reads were trimmed to remove possible adapter sequences and nucleotides with poor quality using Trimmomatic v.0.36. The trimmed reads were mapped to the Homo sapiens GRCh38 with ERCC genes reference genome available on ENSEMBL using the STAR aligner v.2.5.2b. Unique gene hit counts were calculated by using featureCounts from the Subread package v.1.5.2. After extraction, the gene hit counts were assessed for differential expression analysis using DESeq2. The Wald test was used to generate p-values and log2 fold changes. Genes with an adjusted p-value < 0.05 and absolute log2 fold change > 1 were called as differentially expressed genes for each comparison. A gene ontology analysis was performed on the statistically significant set of genes by implementing the software GeneSCF v.1.1-p2. The goa_human GO list was used to cluster the set of genes based on their biological processes and determine their statistical significance.

### Bioinformatic Analyses

Differentially expressed gene (DEG) set filtered by q < 0.01 were used for gene set enrichment analysis (GSEA). Kyoto Encyclopedia of Genes and Genome (KEGG) pathway enrichment analysis was performed in Web Gestalt (Lio and colleagues 2019, http://www.webgestalt.org/). Alternative splicing was analyzed in rMATS. Correlation analysis obtained from DepMap and CCLE.

### In Vitro Proliferation Studies

Procedure was performed as described in (16). In brief, on Day 0, cancer cells were seeded in 96- or 384-well plates. On Day 1, test compounds were added using a Tecan dispenser, and plates were incubated for 7 days. On Day 7, the media was removed and a 1:3 dilution of CellTiter-Glo 2.0 (Promega #G9241) was added. The luminescent signal was then detected by an Envision plate reader. For data analysis, the data were plotted as a percentage of the DMSO control and fitted to the 4-parameter logistic (4-PL) Hill equation using GraphPad Prism.

### MTA and SAM quantification

Procedure was performed as described in (16). In brief, cells were seeded in triplicate in 6-well plates. At indicated time points, one well was used to assess cell count, while extracellular media and intracellular samples were collected from the remaining two wells and stored at -80°C. Intracellular samples were prepared by scraping cells in ice cold 80% MeOH, followed by a vortex, centrifuge, and supernatant transfer for LC-MS/MS analysis. The resulting MTA and SAM data were normalized to their respective internal controls (d3-MTA and 13C5-SAM) using a standard curve for absolute quantification.

### Xenograft Models

Procedure was performed as described in (16). All the procedures related to animal handling, care, and the treatment in this study were performed according to guidelines approved by the Institutional Animal Care and Use Committee (IACUC) of Pharmaron following the guidance of the Association for Assessment and Accreditation of Laboratory Animal Care (AAALAC). In brief, after acclimatization, LN-18 MTAP-deleted tumor cells were injected into mice to form palpable tumors (200-300 mm3). Animals were then randomized to treatment groups and dosed with TNG908. Tumor volume was measured with calipers and analyzed using %TGI and %TV. Statistical comparisons were made with a Two-way ANOVA followed by Dunnett’s or Tukey’s test. Mice were euthanized upon excessive body weight loss or moribundity, and survival analysis was performed using a Kaplan Meier plot and Log-rank test. PK parameters were analyzed using noncompartmental methods by WinNonlin software.

## RESULTS

### TNG908 pharmacodynamic effects are histology agnostic

The recognition of the inherent vulnerability created by MTA accumulation and its impact on PRMT5 activity in *MTAP*-deleted cells has sparked considerable interest in developing novel therapies that exploits this weakness (1, 2). In this context, we have designed a series of MTA-cooperative PRMT5 inhibitors, beginning with TNG908, a highly potent, MTAP-selective, and brain-penetrant compound for the treatment of cancers with *MTAP* deletion (16, 17). Given the variability in PRMT5 inhibitor sensitivity across *MTAP*-deleted tumors and the limitations of protein-based pharmacodynamic markers like SDMA, we sought to identify additional biomarkers that could enhance patient stratification and better capture the functional effects of MTA-cooperative PRMT5 inhibition. To characterize the susceptibility of tumors to PRMT5 inhibition, we conducted a 7-day in vitro viability assay using TNG908 across a curated panel of 29 *MTAP*-deleted cell lines. This panel was intentionally selected to include both previously characterized sensitive and insensitive models, representing tumor types such as melanoma (MEL), pancreatic adenocarcinoma (PDAC), mesothelioma (MESO), lung adenocarcinoma (LUAD), cholangiocarcinoma (CCA), liver hepatocellular carcinoma (HCC), and glioblastoma (GBM). Five *MTAP* wild type cell lines, identified by genomic status within DepMap and confirmed by detectable MTAP expression, were included from PDAC and LUAD lineages, screened under identical conditions, and served as controls. Cell line sensitivity and compound potency were calculated based on the absolute maximal effect (%Amax) derived from the full dose–response curve for each cell line. Values ranged from - 90% to -20% for the *MTAP*-deleted cells and -50% to 20% for the *MTAP*-WT control cells (Fig. 1A). No tumor lineages demonstrated enhanced sensitivity to compound indicating that TNG908 viability effects are lineage agnostic.

**Figure 1.**
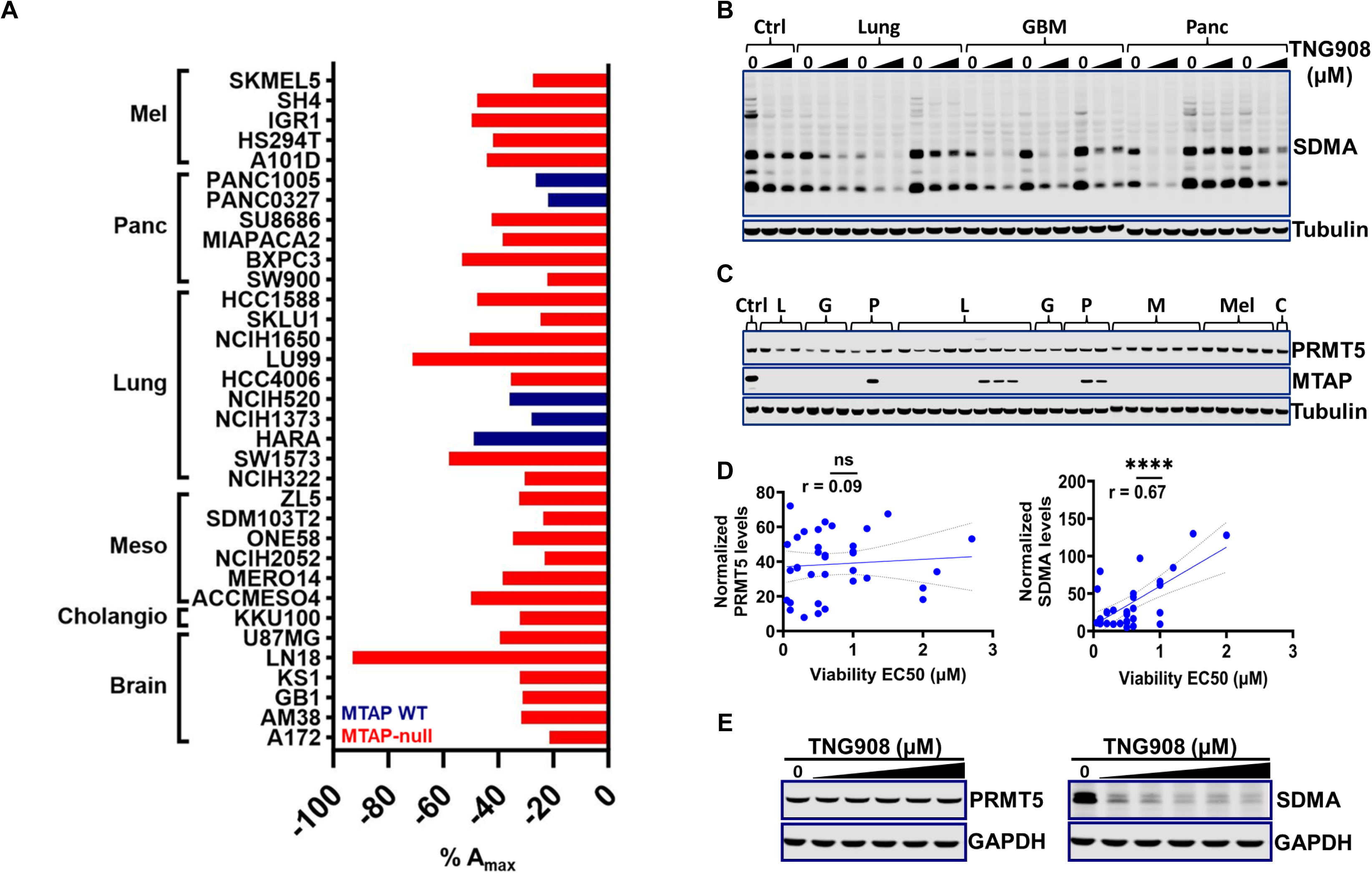
TNG908 pharmacodynamic effects are histology agnostic. **A:** Determination of maximum effect (Amax) in 34 cancer cell lines representing multiple cancer lineages including melanoma, pancreatic adenocarcinoma, mesothelioma, non-small cell carcinoma, cholangiocarcinoma, and glioblastoma following 7-days TNG908 treatment. Cell lines are colored by MTAP status. **B:** Exemplar SDMA immunoblot following a 3-day treatment with TNG908 at cell line-specific EC_20_ and EC_50_ concentrations from (Figure 1A). **C:** PRMT5 and MTAP levels of cell lines from (Figure 1A). **D:** Correlation of normalized PRMT5 (left) or a single SDMA-modified substrate (right) immunoblot levels at the cell line-specific EC_50_. **E:** PRMT5 and SDMA immunoblots following treatment a 3-day treatment of TNG908 at 0.02, 0.08, 0.31, 1.25, 5µM in *MTAP*-deleted LN18 cells.

PRMT5 is the primary type II methyltransferase responsible for depositing SDMA post-translational modifications to downstream substrates and can be used as a global pharmacodynamic marker of PRMT5 activity (4). To confirm the effectiveness of TNG908 in inhibiting PRMT5 activity, we treated the cell panel with cell line specific EC_50_ and EC_20_ concentrations for 3-days and measured the levels of SDMA via western blot. We observed a significant reduction in SDMA modifications in all 29 *MTAP*-deleted cell lines when treated at their respective EC_50_ concentrations, and a more modest but detectable reduction in the 5 *MTAP*-WT cell lines (Fig. 1B, Suppl. 1A). Evaluation of the PRMT5 target SmB/B′, for all 34 cell lines tested demonstrated a significant correlation between reductions in SDMA modifications and viability EC_50_ (Pearson > 0.67) (Fig. 1B, Fig. 1D). Although the viability and SDMA reductions we observed appeared to be broadly lineage agnostic, several cell lines exhibited divergent responses. For example, the GBM cell line A172 displayed minimal loss in viability despite marked SDMA reduction, suggesting on-target engagement without a strong antiproliferative effect. In contrast, the LUSC cell line HC1588 showed robust viability loss with only modest SDMA reduction, indicating a disconnect between the pharmacodynamic marker and phenotypic outcome. These discrepancies suggest that additional, context-dependent factors may influence the cellular response to PRMT5 inhibition. To investigate whether these variances could be clarified by changes in the endogenous levels of PRMT5 or MTAP, western blots were performed (Fig. 1C). As expected, MTAP loss was confirmed in all relevant cell lines, but PRMT5 expression did not correlate with EC_50_ values (Pearson > 0.09) (Fig. 1C, Fig. 1D). PRMT5 protein levels remained unchanged following incubation with 5 µM TNG908, a concentration well above the compound’s in vitro EC_50_ of 110 nM and its established 15-fold selectivity for *MTAP*-deleted cells, indicating that TNG908 does not alter PRMT5 abundance even at suprapharmacologic doses (16, 17). As we have previously reported, TNG908 functions as a potent enzymatic inhibitor of PRMT5 (16, 17). Accordingly, the absence of an effect on PRMT5 protein levels confirms that the observed SDMA reduction is a direct consequence of PRMT5 catalytic inhibition (Fig. 1E). Taken together, these results confirm that TNG908 is a potent and selective MTA-cooperative PRMT5 inhibitor, with functional and pharmacodynamic effects that are dependent on *MTAP* loss and SDMA modulation, and broadly consistent across tumor lineages.

Since *MTAP* deletion leads to accumulation of MTA, resulting in partial inhibition of PRMT5, cells with *MTAP* loss may exhibit baseline aberrations in RNA splicing. Given that PRMT5 inhibition further increases alternative splicing events (ASEs) across multiple cell lines and tumor types (11–14, 18), we hypothesized that the basal AS profiles of untreated cell lines could serve as predictive biomarkers of sensitivity to TNG908. To assess baseline AS patterns, we analyzed RNA-seq data from 29 of the 34 evaluated cell lines obtained from the Cancer Cell Line Encyclopedia (CCLE) (19). Differences in alternative splicing were quantified using percent-spliced-in (PSI) values, representing the ratio of reads including an exon to total reads spanning its junctions. Our analysis specifically focused on a curated subset of functionally relevant exons. Overall, no correlation was observed between TNG908 potency (EC_50_) and the total number of baseline alternative splicing events across cell lines (Fig. 2A) (Pearson > 0.25, Suppl. 2A). This lack of association also held when events were analyzed by subtype, including alternative 3′ splice site (A3SS), alternative 5′ splice site (A5SS), retained introns (RI), and mutually exclusive exons (MXE) (Fig. 2A, Fig. 2B) (Pearson > 0.15–0.28, Suppl. 2B). Given this, we next asked whether a specific subset of splicing events might nonetheless represent a biologically meaningful feature of PRMT5 dependency. To test this, we focused on skipped exon (SE) events as a candidate class for further investigation.

**Figure 2.**
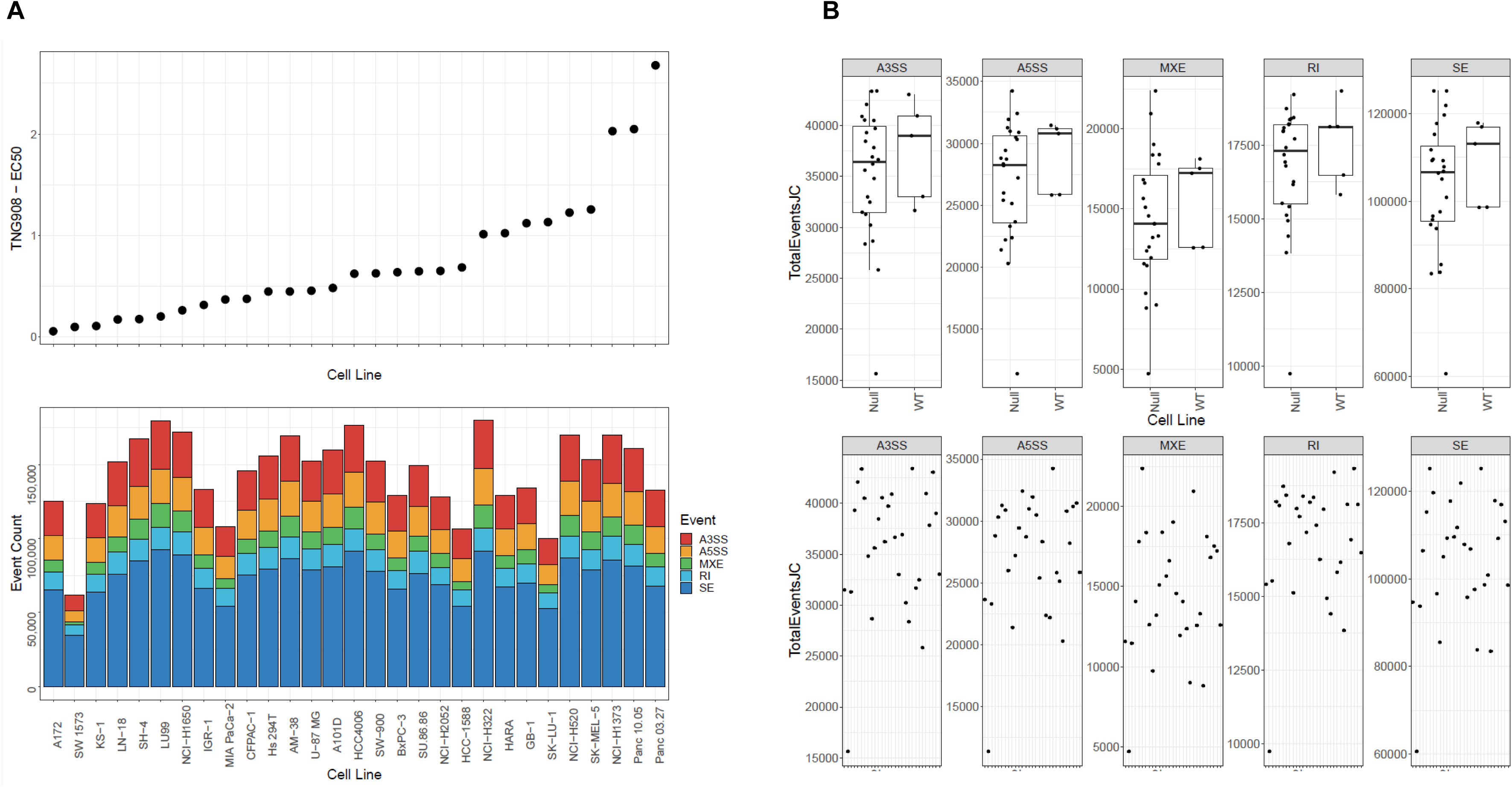
Endogenous alternative splicing does not predict TNG908 sensitivity. **A:** Assessment of total event counts as determined by PSI expression from RNA-seq data extracted from CCLE for 29 cancer cell lines from various cancer lineages. Cell lines ranked by TNG908 EC_50_ values (Top). Bars separated by specific event type Alternative 3’SS (A3SS), Alternative 5’SS (A5SS), Mutually exclusive exons (MXE), Retained introns (RI) and Skipped exons (SE) (Bottom). **B:** Events, as measured by junction counts, separated by class were totaled, averaged, and compared in both *MTAP*-deleted and *MTAP*-WT cells (Top). Total junction counts per individual cell lines (Bottom).

### PRMT5 inhibition induces alterations in alternative splicing

Proper spliceosome function is partially dependent on PRMT5 activity, which plays a critical role in early spliceosome assembly. Specifically, PRMT5 methylates the arginine/glycine-rich C-terminal tails of Sm proteins (SmB/B’, SmD1, SmD3), enhancing their affinity for the survival motor neuron (SMN) complex. This post-translational modification facilitates snRNA binding, maturation, and nuclear import, enabling efficient spliceosome assembly (20).

Although no baseline splicing category correlated with TNG908 sensitivity, we reasoned that focusing on a specific subset of events might reveal a biologically meaningful feature of PRMT5 dependency. We therefore selected SE events for further investigation, given their potential to directly disrupt protein function and their recurrent modulation following PRMT5 inhibition. To do this, we performed bulk RNA-seq on *MTAP*-deleted cell lines representing GBM (LN18), NSCLC (A549), and PDAC (Miapaca2) following a 3-day treatment with TNG908 at each cell line’s EC_50_. Across the three cell lines, we identified 5,467 statistically significant differentially spliced events, with the majority being RI (15%, 820) and SE (62%, 3,401) events (Fig. 3A). These findings are in line with prior reports and indicate that PRMT5 inhibition produces widespread changes in alternative splicing, with a predominance of SE events among affected splicing types (11–14, 18). Among the SE events, 238 (4.3%) were shared across all three *MTAP*-deleted cell lines (Fig. 3B), suggesting that a subset of alternative splicing changes reproducibly occurs following TNG908 treatment regardless of tumor type. Gene set enrichment analysis (GSEA) of the AS event list revealed enrichment for genes involved in RNA processing, cell-cycle regulation, and DNA damage (Fig. 3C).

**Figure 3.**
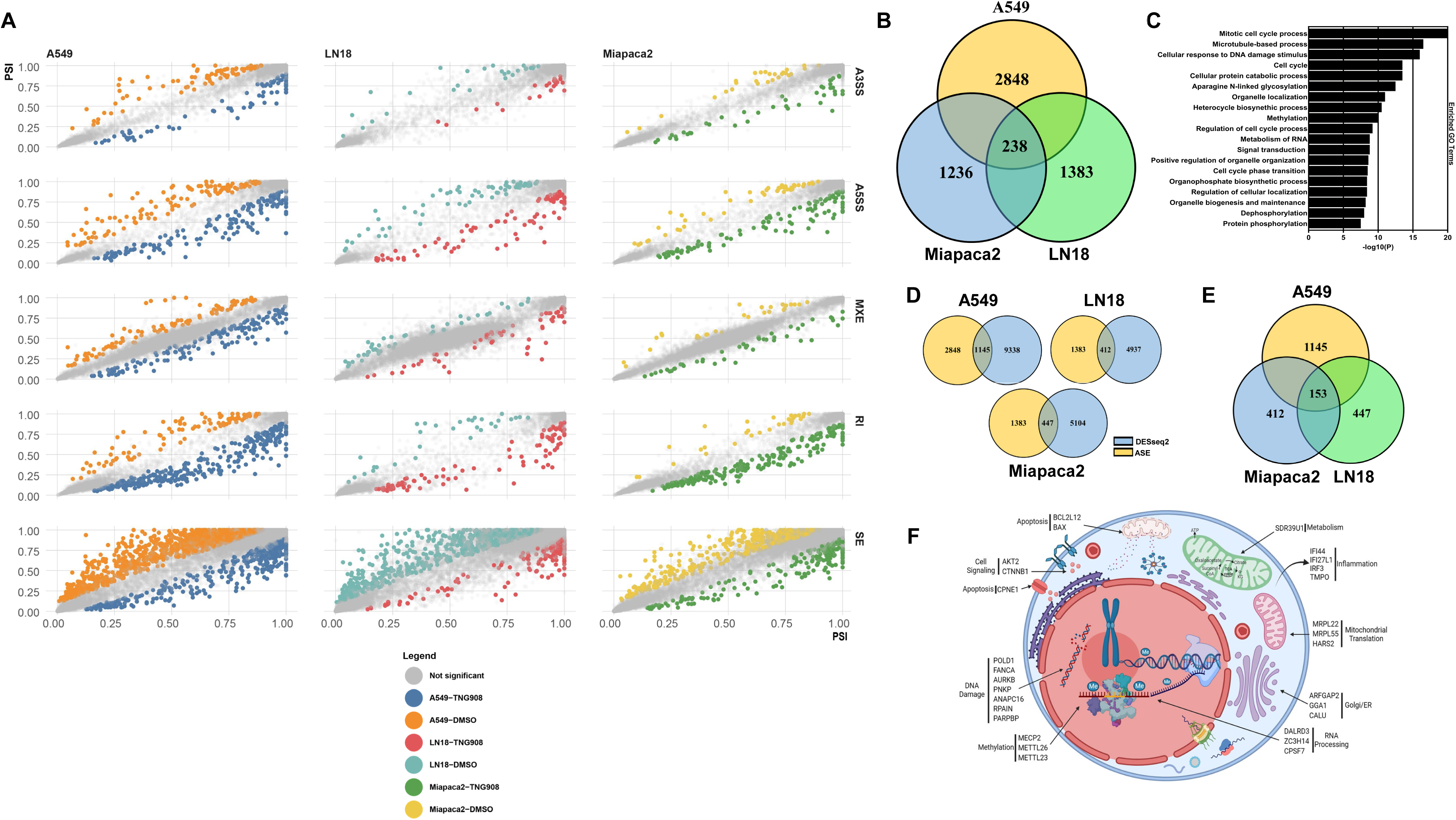
PRMT5 inhibition induces alterations in alternative splicing. **A:** rMATS (replicate Multivariate Analysis of Transcript Splicing) analysis was performed on the RNA-seq data from 3-day TNG908 treatment of LN18, A549 and Miapaca2 cells to detect differential alternative RNA splicing. **B:** Venn diagram of identified alternative events between cell lines. **C:** Gene ontology analysis of significantly regulated genes from (Figure 2). **D:** Venn diagrams of the differential gene expression from the *MTAP*-deleted cell lines treated for 3 days with 1μM TNG908 and the identified alternative splicing events. **E:** Venn diagram of the alternatively spliced associated differentially expressed genes in all *MTAP*-deleted cell lines. **F:** Diagram demonstrating function of a subset of genes with validated alternative events.

To examine the relationship between alternative splicing and gene expression, we filtered RNA-seq data to remove genes with low or no expression, then performed differential expression analysis using DESeq2 (21) across all three cell lines. While transcript abundance was largely independent of alternative splicing status, genes with splicing changes were more likely than others to also show altered expression (Fig. 3D). We next identified the intersection of these two groups, differentially expressed genes that also exhibited alternative splicing consistently in all three cell lines yielding a set of 153 genes (Fig. 3E). Collectively, these findings indicate that PRMT5 inhibition has a reproducible impact on a specific subset of genes at both the splicing and expression levels, many of which are involved in RNA processing and related pathways.

To validate the RNA-seq identified SE events induced by PRMT5 inhibition, we conducted splicing assays using primers designed to amplify either the canonical transcript or the alternative splice isoform identified in Figure 3. From the set of 153 genes that exhibited both differential expression and alternative splicing in all three cell lines (Fig. 3E), we selected 40 SE events (PSI change > 20%) for validation and confirmed 36 (90% validation rate). The catalog of validated events, selected based on the level of PSI, consistency across cell lines, and accurate read mapping, includes genes with diverse biological functions such as RNA processing, apoptosis, DNA damage response, methylation, and inflammation (Fig. 3F). The breadth of these functional categories indicates that alternative splicing changes following PRMT5 inhibition could contribute to multiple downstream cellular effects, in line with the varied biological roles previously associated with PRMT5.

### Identified splicing alterations are dose responsive and histology agnostic

We next questioned whether the identified splicing events are a direct and specific consequence of PRMT5 inhibition and whether this effect is selective to *MTAP*-deleted cells. To quantify and characterize the relationship between TNG908-induced PRMT5 inhibition and the level of SE response, dose-response curves were generated for 22 events. We extracted RNA from LN-18 *MTAP*-deleted cells following a 3-day incubation with TNG908 dosed from 0.001-10 µM. This dose range was chosen to span the full spectrum of TNG908 activity, as the compound has an EC_50_ of approximately 0.1 µM in this cell line (Suppl. 3A). Analysis of event-specific PSI for each product established that several events were sensitive to PRMT5 inhibition in a dose-dependent fashion, though this effect was not observed across all events (Fig. 4A). Plotting the IC_50_ for each splicing event revealed distinct drug response profiles, suggesting that individual splicing defects vary in their sensitivity to PRMT5 inhibition (Fig. 4A). To confirm the selectivity of the identified events, we focused on the subset that showed a clear dose-responsive relationship to TNG908 treatment in LN18 *MTAP*-deleted cells (Fig. 4A). We then compared these events in an isogenic model generated by reintroducing MTAP into the endogenous *MTAP*-deleted GBM line LN18, as reported previously (16, 17). *MTAP*-WT and *MTAP*-deleted LN18 cells were treated with 1μM TNG908 for 3 days, RNA was extracted, and PSI levels were quantified. Analysis revealed minimal splicing alterations across these events in *MTAP*-WT cells, in contrast to the pronounced changes observed in *MTAP*-deleted cells, confirming that the splicing effects of TNG908 are selective to the *MTAP*-deleted background (Fig. 4B).

**Figure 4.**
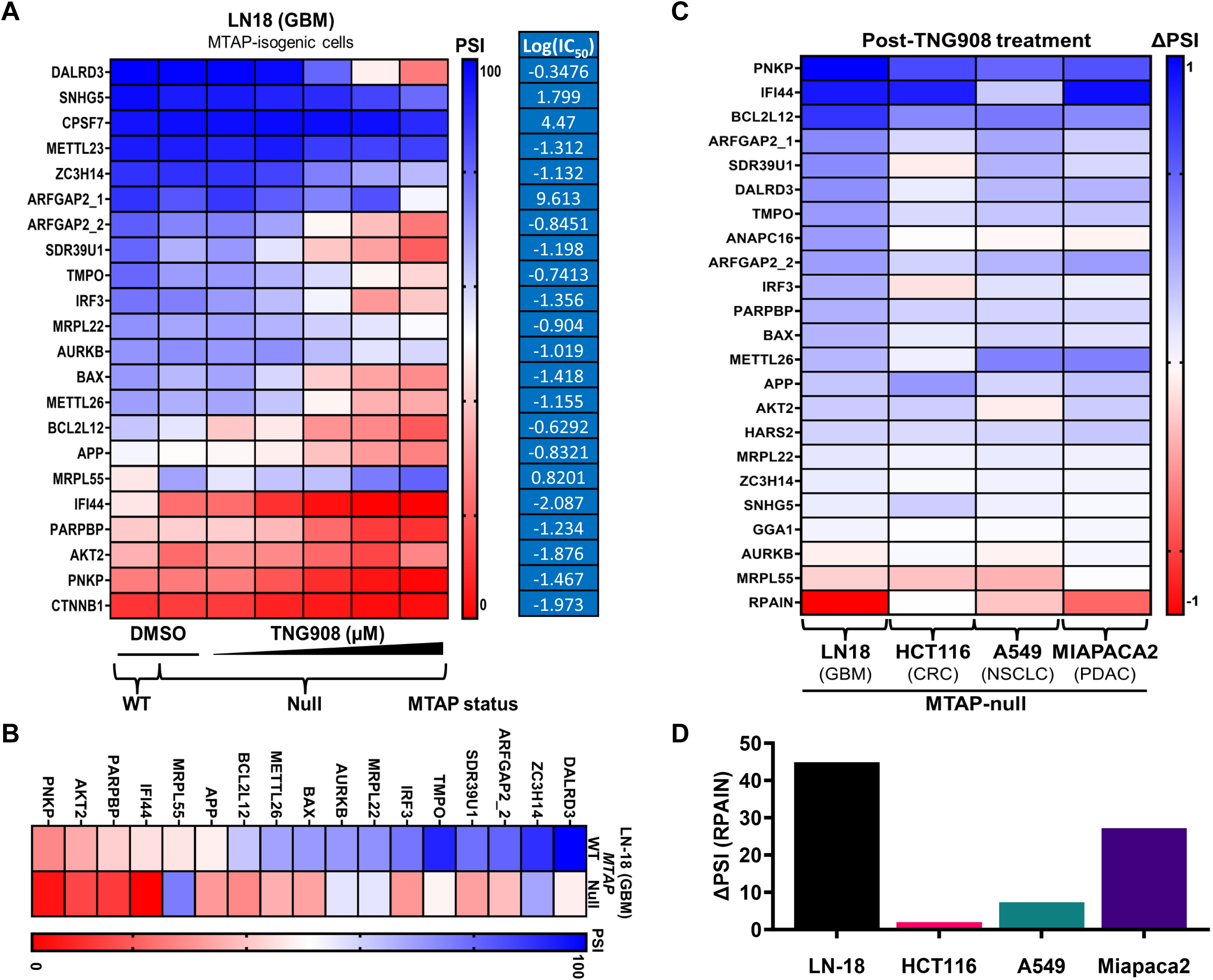
Validated splicing alterations are dose responsive and histology agnostic. **A:** Heatmap of 22 distinct alternative splicing events (ASE) following 3-days TNG908 treatment in the *MTAP*-deleted GBM LN-18 cell line. Data reported as percent spliced in (PSI) of the alternative exons (% inclusion = inclusion/sum of inclusion + exclusion). Table of PSI values transformed, plotted, and expressed as LogIC_50_ values. **B:** Heatmap of 22 ASEs expressed as PSI following 3-day treatment of 1µM TNG908 in LN-18 *MTAP*-WT or LN-18 *MTAP*-deleted cells. **C:** Heatmap of 22 distinct ASEs from *MTAP*-deleted cells treated for 3-days with treated with 1µM TNG908, the maximally efficacious dose identified from Figure 2.1. Data reported as ΔPSI of the alternative exons (ΔPSI = PSI DMSO – PSI TNG908). Cell lines represent glioblastoma (GBM), colorectal cancer (CRC), non-small cell lung cancer (NSCLC), and pancreatic adenocarcinoma (PDAC). **D:** ΔPSI of RPAIN ASE for treated cell lines.

Alternative splicing is a post-transcriptional mechanism of gene regulation that generates distinct isoforms in different tissues, altering protein function, localization, and activity. This is due to a variety of factors including the recognition of weak and strong splice sites, presence of *cis*-regulatory elements and splicing factor expression levels (20). Thus, the formation of a specific alternatively spliced isoform could be context dependent. Given that MTAP loss occurs in ∼10–15% of cancers across diverse tumor types, we questioned whether the validated splicing events occur in *MTAP*-deleted cancer cells regardless of cellular histology. We isolated RNA from *MTAP*-deleted cell lines representing GBM, PDAC, non-small cell lung cancer (NSLC) and colorectal cancer (CRC) lineages following a 3-day 1µM dose of TNG908. The relative abundance of each of the 22 splice events were compared to their isoform and cell line specific DMSO control and expressed as ΔPSI to quantify and compare changes in splicing patterns (Fig. 4C). Analysis of ΔPSI for each event revealed that although we were able to detect the predicted alternative isoform in each *MTAP*-deleted cell line, splicing events are not perturbed uniformly relative to each line. For example, the level of alteration for RPA-interacting protein (RPAIN) isoform is higher in LN18 cells compared to the other lines (Fig. 4D). This observation may not be surprising, as each cell line was not dosed at its cell-line specific EC_50_, and each line varies in its relative sensitivity to PRMT5 inhibition (Suppl. 3A, 3B). Taken together these data demonstrate that the identified alternative events can be further mediated by PRMT5 inhibition and are not specific to tumor lineage.

### Alternative splicing events can be specifically induced by PRMT5 inhibition

Alternative splicing can be triggered by various factors, including treatment with cytotoxic agents, disruptions to spliceosome components, and cellular insults leading to DNA damage (22). Splicing can also be modulated indirectly, as seen with a recent study detailing the interplay between CDK4/6 inhibition and MEP50:PRMT5 activity (23, 24). Therefore, to determine whether the identified events are specifically and directly caused by PRMT5 inhibition, we conducted splicing assays in four different *MTAP*-deleted cancer cells representing multiple tumor histologies using the SE event DALRD3, as identified in Figure 4A. DALDR3 was chosen as an example due to the sensitivity and dynamic range of the SE splicing event. To determine whether the reduction of PRMT5 activity, either directly or indirectly, affects the splicing of DALRD3, we utilized two inhibitors: TNG908, a direct PRMT5 inhibitor, and AG-270, an inhibitor of MAT2A that functions upstream to deplete the PRMT5 substrate, S-adenosyl methionine (SAM). MAT2A is responsible for catalyzing the synthesis of the MTA precursor SAM from methionine and ATP and is the subject of multiple drug discovery efforts (25). To explore whether the observed alternative splicing events were facilitated by spliceosome perturbation or cell cycle regulation, we employed the SF3B1 inhibitor herboxidiene, the CDK4/6 inhibitor palbociclib, and the microtubule destabilizer docetaxel, which induces G2/M cell cycle arrest. To ensure that the observed splicing phenotypes were not a secondary consequence of cytotoxicity, each compound was dosed below their respective IC_50_ concentration which significantly impacts cell viability in LN-18 *MTAP*-deleted cells (Suppl. 4A). Following a 3-day treatment RNA was isolated and the PSI for DALRD3 was compared for each treatment (Fig. 5A). We observed that the splicing of DALRD3 is only discernible following the inhibition of PRMT5. To validate this discovery, we performed splicing assays on several other events and found that while most of the identified events were consistently regulated by PRMT5 inhibition, some events like ARFGAP2 were also sensitive to other splicing perturbants (Fig. 5B). To distinguish between splicing events caused by drug activity versus those resulting from a general reduction in cellular health, we conducted kinetic splicing assays in LN-18 *MTAP*-deleted cells using TNG908 at cell line specific EC_90_ (1µM) and EC_25_ (0.01µM) concentrations. RNA was extracted after 1, 3 and 5 days and PSI was calculated for the splicing events DALRD3 and PNKP. Analysis of PSI for each event demonstrates that splicing is initiated at 1 day and is maintained for up to 5 days (Fig. 5C). A comparison of the kinetics of DALRD3 splicing and Caspase 3/7 induction revealed that splicing is initiated at an earlier time point than PRMT5-induced apoptosis (Fig. 5B, Fig 5D).

**Figure 5.**
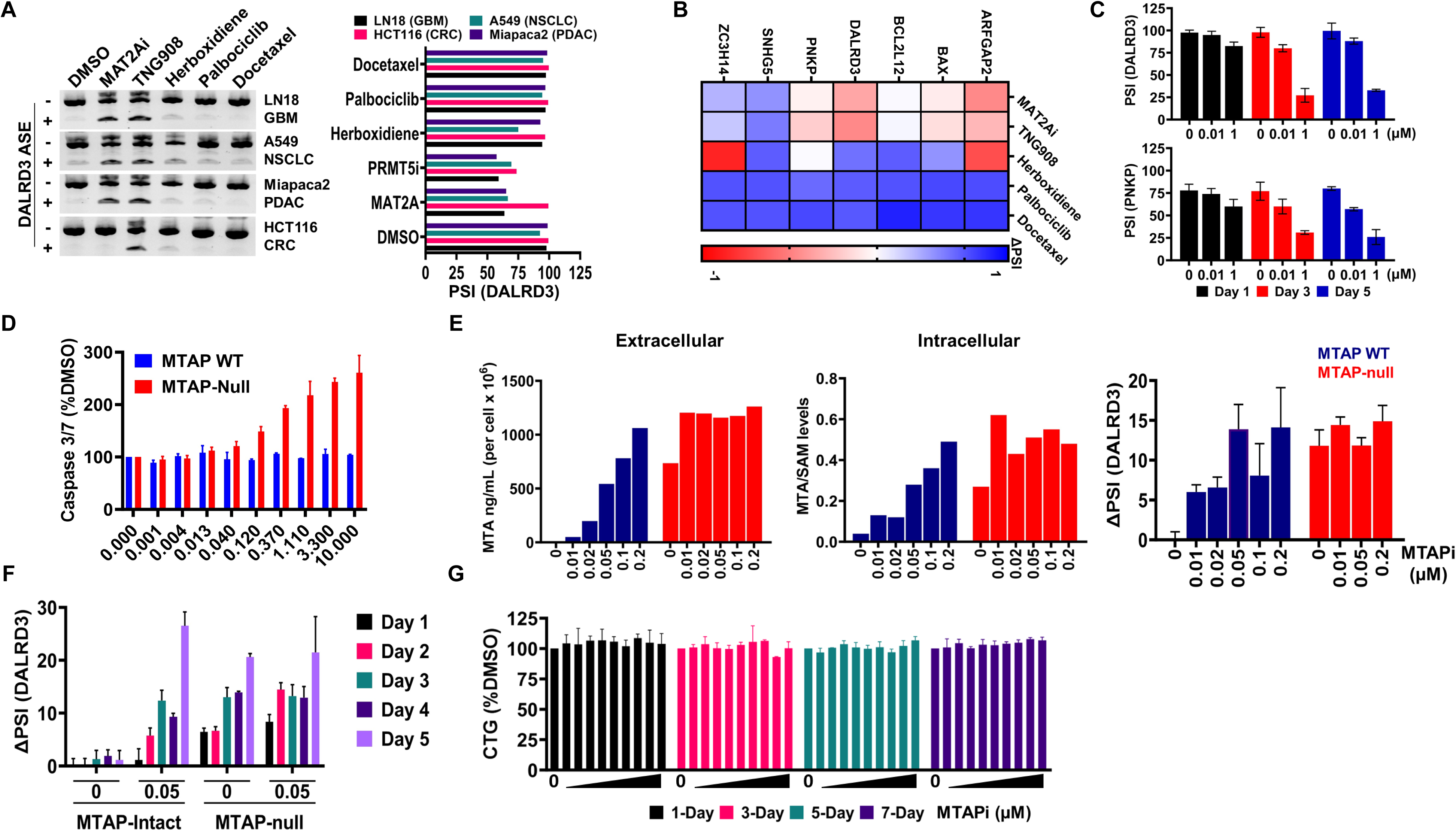
Subset of validated events are specific for PRMT5 inhibition. **A:** DALRD3 ASE raw data (left) or PSI (right) following a 3-day treatment of indicated compounds dosed at their IC_50_ concentrations in *MTAP*-deleted cell lines representing the indicated histologies (LN18, GBM; HCT116, CRC; A549, NSCLC; Miapaca2, PDAC). MAT2Ai is AG-270. **B:** Heatmap of ΔPSI (ΔPSI = PSI DMSO – PSI TNG908) values from ARFGAP2, BAX, BCL2L12, DALRD3, PNKP, SNHG5 and ZC3H14 ASEs following TNG908 treatment for 3 days in LN18 *MTAP*-deleted cells. **C:** PSI of DALRD3 and PNKP ASEs following TNG908 treatment for indicated days in LN18. Data are presented as mean ± SD. **D:** Caspase 3/7 activity in LN-18 *MTAP*-deleted cells following a 3-day treatment of TNG908. **E:** (Left and middle panel) LC-MS/MS analysis of MTA and SAM metabolite levels following MTAP inhibitor (MTDIA) treatment for 3 days. (Right) ΔPSI of DALRD3 SE (ΔPSI = PSI_MTAPi_ - PSI_MTAP WT_ DMSO) following MTAPi treatment for 3 days at indicated doses in the HAP1 *MTAP*-isogenic cell line pair. Data are presented as mean ± SD. **F:** ΔPSI of DALRD3 SE (ΔPSI = PSI_MTAPi_ - PSI_MTAP WT_ DMSO) following MTAPi treatment at indicated doses and times in the HAP1 *MTAP*-isogenic cell line pair. Data are presented as mean ± SD. **G:** Antiproliferative activity of MTAPi in HAP1 *MTAP*-WT cells (10µM top dose) for indicated time points in CellTiter-Glo activity assay. Data are presented as mean ± SD.

Both our research and other studies have demonstrated that *MTAP*-deleted tumor cells accumulate intracellular MTA, which can also be monitored in extracellular cell culture media (1–3). These elevated levels contribute to the selective sensitivity of MTA-cooperative PRMT5 inhibitors and might predict a response to drug treatment. While liquid chromatography-tandem mass spectrometry (LC-MS/MS) is the primary method for evaluating metabolite levels like MTA, its application can present technical challenges and may not be standardized for routine clinical use. To formally demonstrate that MTA is the causal driver of the splicing alterations observed in *MTAP*-deleted cells, we hypothesized that the accumulation of MTA in an *MTAP-WT* cell would induce the same splicing changes we observed in the *MTAP*-deleted cells, thereby providing a formal link between MTA-driven PRMT5 inhibition and these specific splicing events. To achieve this, we utilized the potent MTAP inhibitor, MTDIA (26, 27), in a HAP1 MTAP isogenic cell line pair to directly mimic the biochemical state of an *MTAP*-deleted cell. The drug was administered at a dose that led to an intracellular MTA/SAM ratio comparable to that found in the *MTAP*-deleted line (Fig. 5E). To confirm this, we quantified the absolute amount of MTA in both cell culture media and cell lysates via LC-MS/MS. Inhibition of PRMT5 has no impact on the intracellular levels of SAM as quantified, thus SAM was used as a normalization factor to determine the intracellular levels of MTA. Analysis of the resolved peaks revealed that MTA levels exhibited a proportional increase in response to the MTDIA dose in the *MTAP*-WT cells. A slight increase was also observed in the *MTAP*-deleted cells which could be attributed to low levels of MTAP found in the cell culture media (28). In a similar fashion to the metabolite analysis, RNA was extracted from a replicate plate and the presence of the DALRD3 AS event was compared to the increased levels of MTA in the *MTAP*-WT line. At 50nM, MTDIA increased intracellular MTA to levels nearly identical to the basal concentration found in *MTAP*-deleted cells (Fig. 5E). Remarkably, at this same dose, the DALRD3 splicing event was induced to a comparable extent as in the *MTAP*-deleted line, demonstrating a tight correlation between MTA accumulation and functional PRMT5 inhibition. This relationship was further reinforced in a kinetic analysis where 50nM MTDIA triggered DALRD3 splicing in *MTAP*-WT cells within just two days, and the response was maintained for at least five days (Fig. 5F). These findings not only confirm MTA as the causal driver of specific splicing alterations but also highlight DALRD3 as a sensitive and durable functional readout of PRMT5 inhibition driven by MTAP loss. To determine if the inhibition of MTAP causes alterations to cellular fitness we evaluated the viability effects of MTDIA dosed cells and found no impact to cellular health after 1, 3, 5 and 7-days (Fig. 5G). Administration of exogenous MTA reduced SDMA-modified protein levels in *MTAP*-WT cells within one day (Suppl. 4B), confirming that MTA alone is sufficient to inhibit PRMT5 and drive the observed splicing changes, further reinforcing the role of MTA as the driver of the observed splicing alterations. Taken together we demonstrate that the identified events are specifically dependent on PRMT5 either through direct pharmacological intervention or by the accumulation of MTA.

### Alternative splicing events correlate with efficacy in vitro

Considering that MTA accumulation induces alternative splicing, we hypothesized that cell lines with enhanced alternative splicing may exhibit greater sensitivity to an MTA-cooperative PRMT5 inhibitor due to elevated levels of MTA. Therefore, our next inquiry focused on whether the extent of the PRMT5 dependent SE events could serve as a determinant of TNG908 sensitivity within the characterized cell line panel. We utilized all 34 cell lines from the panel (Fig. 1A) and treated each cell with their cell line specific EC_50_ concentrations or DMSO for 3 days. RNA was extracted and all cell lines were simultaneously run on a large 2% acrylamide gel to directly compare the splicing levels of a particular event. We first evaluated the basal splicing levels of each cell line for 10 PRMT5 dependent splicing events. As anticipated, substantial differences in the amount of PSI were observed when comparing the untreated cell lines. However, by ranking the cells in order of their EC_50_ as determined by a 7-day viability assay no clear trend was observed with any AS event (Fig. 6A). This result agrees with the data we obtained from our CCLE analysis where no direct trend was observed. We next evaluated the ΔPSI of each cell line to compare the effect of TNG908 treatment on the AS for each event in all cell lines. We observed a strong induction of the alternative events in all cell lines tested irrespective of their sensitivity to PRMT5 inhibition (Fig. 6B). This result is not unexpected since each cell line was dosed with a concentration of TNG908 capable of inducing a half-maximal viability effect. These data also imply that the levels of MTA required for TNG908 to cooperatively bind to PRMT5 are sufficient for the amount of inhibitor used for each cell line. These data suggest that the potency of alternative splicing inhibition is a robust predictor of the cytotoxic potency of TNG908.

**Figure 6.**
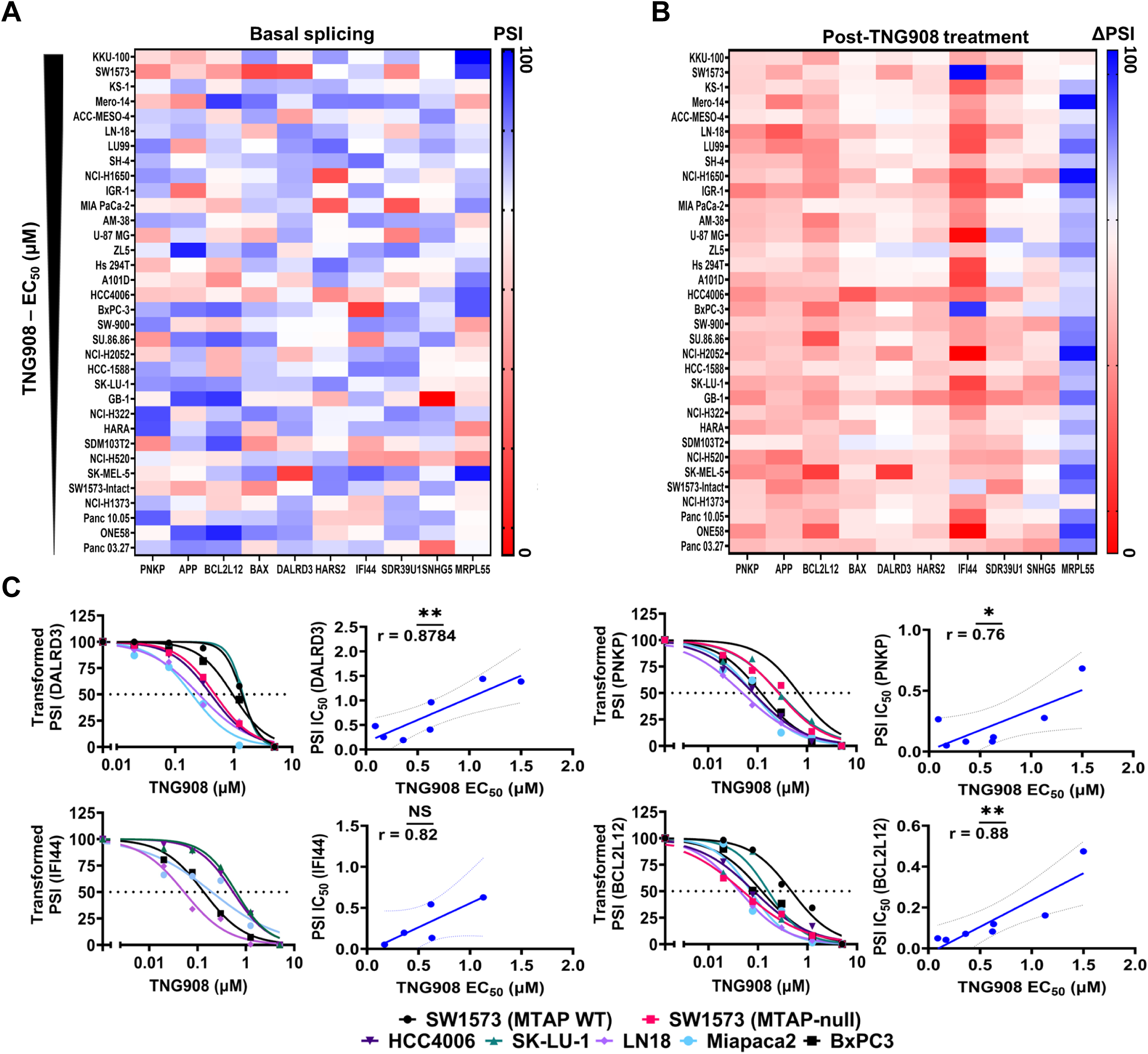
Alternative splicing events correlate with efficacy, but do not predict sensitivity to TNG908. **A:** Heatmap of basal alternative splicing in cell lines from Figure 3A ranked by sensitivity toTNG908. **B:** Heatmap of ΔPSI of cell lines treated with TNG908 for 3 days at their specific EC_50_. **C:** PSI of ASEs following 3-day TNG908 treatment in the indicated cell lines. PSI values transformed and linear regression plots were constrained at top and bottom (left). Correlation of PSI IC_50_ and viability EC_50_ (right).

Since all cell lines were dosed at a concentration capable of inducing the same degree of cell viability loss we questioned whether the PSI-dependent IC_50_, obtained in a subset of cell lines, would correlate with PRMT5-dependent efficacy. We therefore took 6 *MTAP*-deleted cell lines representing LUAD, PDAC and GBM lineages and a reconstituted *MTAP*-WT cell line to use as a control. Each cell line was dosed for 3 days with an ascending dose of TNG908 (0.02-5µM), and calculated PSI values were compared to the cell line specific DMSO control. The subsequent PSI expression values were transformed and plotted using linear regression and the PSI IC_50_ for each cell was compared. For the DALRD3 SE event we observed a strong correlation (Pearson > 0.87) between the response to TNG908 viability and induced alternative splicing across all cell lines, with the LN-18 cell line, on average, being more sensitive to PRMT5 induced AS (Fig. 6C). Similar findings were observed for PNKP SE (Pearson ≥ 0.76), IFI44 SE (Pearson ≥ 0.82) and BCL2L12 SE (Pearson ≥ 0.88) suggesting that alternative splicing may be a potential pharmacodynamic biomarker to evaluate TNG908 efficacy in vitro. The correlation between these splicing events and TNG908 efficacy is stronger and more robust than the correlation observed for SDMA levels (Fig. 1D). This more pronounced relationship, coupled with the potential for a greater dynamic range, suggests that alternative splicing events may serve as a more sensitive pharmacodynamic biomarker for the effective reduction of SDMA by TNG908.

### PRMT5 inhibitor induced alternative splicing events occur in vivo

Since the PRMT5 dependent alternative splicing events can be used to evaluate the on-target efficacy of TNG908 in vitro we questioned whether these events can be identified in vivo and serve as a pharmacodynamic biomarker. In a previous study, we observed that TNG908 treatment drives strong PRMT5 inhibition in a 7-day PK/PD study using the LN18 CDX model (16). To support the relevance of splicing changes as direct pharmacodynamic effects rather than downstream consequences of tumor regression or loss of viability, we conducted a short-term PK/PD study in the LN-18 *MTAP*-deleted CDX model. TNG908 was dosed at 1, 3, 10, 30, and 60 mg/kg BID for four days (n=4 per group), and plasma and tumor samples were collected two hours after the final dose. TNG908 plasma levels ranged from 100 to 10,000 ng/mL across dose groups, and SDMA-modified protein levels for a single PRMT5 substrate were reduced by nearly 75% at 30 and 60 mg/kg (Fig. 7B). As expected, dose-dependent antitumor activity was observed, consistent with previous findings (Fig. 7A) (16).

**Figure 7.**
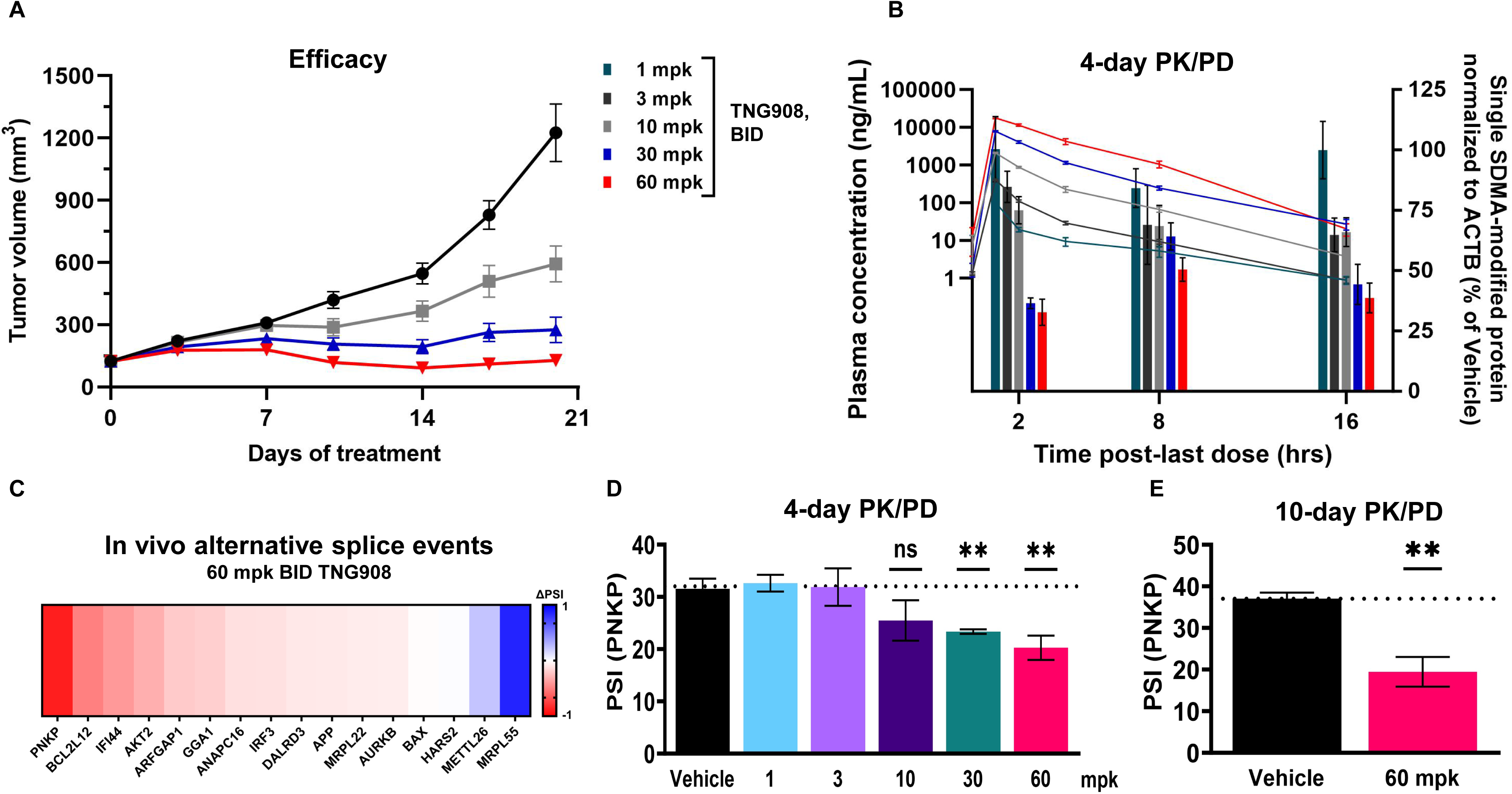
TNG908-induced alternative splicing events occur in vivo. **A:** Efficacy study in the LN18 *MTAP*-deleted subcutaneous GBM CDX model. TNG908 or vehicle was dosed BID, and PK and tumor samples were collected at the indicated timepoints. N=4 tumors per group, data are presented as mean ± SEM. **B:** PK/PD in the LN18 *MTAP*-deleted subcutaneous GBM CDX model. TNG908 or vehicle was dosed BID, and PK and tumor samples were collected at the indicated timepoints. N=4 tumors per group and data are presented as mean ± SEM. **C:** ΔPSI of the alternative exons (ΔPSI = PSIDMSO - PSITNG908) from tumors from (Figure 2.27) extracted 2-hours post last dose. **D:** PSI of PNKP ASE in xenografts following TNG908 treatment for 4 days at indicated doses. Data are presented as mean ± SD from 4 samples. **E:** PSI of PNKP ASE in xenografts following TNG908 treatment for 10 days. Data are presented as mean ± SD from 4 samples.

Given that TNG908 markedly decreases SDMA-modified protein levels in an *MTAP*-deleted CDX model, our next aim was to establish whether the SE events could be detected in vivo. To determine this, we assessed PSI values from purified RNA extracted from terminal tumor samples (n=4) following a 4-day, 60 mg/kg BID dosing regimen of TNG908. To minimize the likelihood that observed splicing events were secondary to tumor regression or cell death, we selected a timepoint characterized by robust SDMA loss with minimal impact on tumor growth. Each PSI value was compared to the vehicle control group and expressed as ΔPSI. We identified several alternative events which could be distinguished in vivo, including the DNA damage modulator PNKP (Fig. 7C). Although the DALRD3 SE event was profiled in vitro due to its sensitivity and dynamic range, no detectable alteration was observed in vivo in the LN18 *MTAP*-deleted CDX model. Since PNKP was a strong initial hit, we next assessed the dose responsive relationship of PNKP alternative splicing and PRMT5 inhibition. Importantly, PSI alterations were not only significantly decreased with increasing TNG908 dose but also appear to be consistent with the SDMA reductions observed in the same study (Fig. 7D). To determine whether the PNKP splicing response is durable and not simply a consequence of tumor regression, we assessed PSI values at a later time point. Tumor samples collected after 10 days of TNG908 treatment (60 mg/kg, n=4) still showed significantly altered PNKP splicing, despite minimal tumor shrinkage at this time point (Fig. 7E, Fig. 7B). These results demonstrate that PRMT5-dependent alternative splicing in vivo is both sustained and uncoupled from gross effects on tumor viability.

## Discussion

The concurrent deletion of MTAP with the tumor suppressor CDKN2A occurs in ∼10–15% of human cancers, and therapeutic targeting of this co-deletion could benefit a patient population with otherwise limited treatment options (29). Targeting PRMT5 by exploiting the heightened MTA accumulation in *MTAP*-deleted tumors represents a novel therapeutic strategy with strong clinical potential. In this study, we show that PRMT5 inhibition with TNG908 consistently reduces growth across 29 *MTAP*-deleted cell lines representing diverse tumor lineages. We further demonstrate that post-translational SDMA modifications are selectively reduced in *MTAP*-deleted versus *MTAP*-WT cells, supporting the broad applicability of MTA-cooperative PRMT5 inhibitors across tumor types. Importantly, correlation analyses identified SDMA reduction, not PRMT5 expression, as the main driver of differential vulnerability, suggesting the magnitude of SDMA reduction could serve as a pharmacodynamic biomarker of early therapeutic efficacy. However, prior clinical trials of PRMT5 inhibitors, both SAM-competitive and SAM-cooperative, have shown robust SDMA suppression without corresponding tumor regression (41–46), indicating that while SDMA reflects target engagement, it may not fully capture therapeutic response.

Currently no arginine demethylase has been identified, suggesting that arginine methyltransferases deposit stable and less-dynamic epigenetic modifications compared to other post-translational modifications (30, 31). Biochemical and cellular assays have also observed long residence time and sustained inhibition of PRMT5 even after compound washout (32). This is important to note since epigenetic alterations generally take longer to induce their desired effect and may explain why even though SDMA modifications are abolished immediately following PRMT5 inhibitor treatment, viability effects lag significantly behind. Our data agrees with this assessment since PRMT5 inhibition in the LN-18 *MTAP*-deleted CDX model required a minimum of 14-days of 60 mg/kg BID dosing to achieve maximal anti-tumor response. In contrast SDMA modifications, the most proximal measurement of PRMT5 inhibition, were significantly reduced following a 4-day drug treatment, supporting our claim that reductions to SDMA are essential for determining functional inhibition but may be inefficient at demonstrating efficacy. Therefore, a companion biomarker that correlates with SDMA and represents a robust, sensitive, and specific pharmacodynamic response could be highly informative in future clinical studies.

PRMT5 has well established pleiotropic effects, encompassing epigenetic regulation to direct substrate inhibition. For this study our focus was on the critical role PRMT5 plays in maintaining RNA processing and splicing fidelity and whether this intrinsic biology could be harnessed to create a pharmacodynamic biomarker. Following a 3-day inhibition we found that TNG908 induces significant gene expression and alternative splicing changes to GBM, NSCLC and PDAC tumor lineages. Consistent with other published reports (11–14), the alterations observed in all tested cell lines exhibited changes in RNA processing, DNA damage, and cell-cycle regulation. Additionally, we observed significant downregulation in the differential expression of several DDR-related proteins, including ATM, PNKP, POLD1, FANCA, and BRCA1/2. Based on these protein expression changes, we hypothesize that PRMT5 inhibition may alter DNA damage response pathways in *MTAP*-deleted tumors, warranting future investigation of potential combination approaches. We also identified a subset of events that exhibited consistency across 29 unique *MTAP*-deleted tumors, indicating that these events may represent a smaller set within a larger core of splicing events. Furthermore, our data indicate that these events are specifically dependent on the activity of PRMT5, either through MTA accumulation or direct inhibition. This implies that the distinctive mechanism of PRMT5 in methylating the Sm proteins may generate a core set of splicing alterations specific to PRMT5 inhibition. Notably, despite hypothesizing that baseline splicing burden would predict sensitivity to PRMT5 inhibition, no correlation was found between endogenous ASE levels and TNG908 potency. This contrasts with a prior report linking intron retention to PRMT5 dependency (10), likely reflecting differences in inhibitors, cell line panels, or sensitivity metrics (e.g., AUC vs. EC_50_).

While baseline splicing was not predictive, PRMT5 inhibition reproducibly induced a consistent panel of alternative events across tumor types, reinforcing their lineage-agnostic nature. These findings suggest that the predictive potential of splicing lies not in overall ASE burden but in the qualitative features of specific event classes. For instance, alterations in SNHG5 were observed in all lines tested, accompanied by significant shifts in expression of other SNHG family members. Given the role of SNHG genes in snoRNA biogenesis and transcriptional regulation (33, 34), and reports linking SNHG5 to reduced EGFR expression (35), it is notable that PRMT5 inhibition here correlated with increased EGFR expression, suggesting that altered SNHG5 isoforms may modulate this pathway.

Other characterized events also highlight potential mechanistic links. DALRD3 and METTL26, both members of the class I methyltransferase superfamily, shifted from their canonical isoforms under PRMT5 inhibition. DALRD3 disruption impairs tRNA m³C formation and causes severe neurologic phenotypes (36), while METTL26 is predicted to play roles in protein modification (37, 38). These findings raise the possibility that altered splicing of these enzymes disrupts tRNA maturation and broad cellular functions. Likewise, the consistent alteration of IFI44 across tumor types, and its known regulation of IRF3 activity (39), aligns with emerging evidence that PRMT5 inhibition enhances IRF3-mediated innate immune signaling (40).

The challenge of defining a splicing-based patient stratification signature likely reflects biological variability. A recent study of GBM samples found only 3% overlap in altered transcripts across lines (14), underscoring the influence of genetic context. We attempted to mitigate this by validating events across a diverse panel, yet variability persisted. Nonetheless, the strong correlation between PSI IC_50_ values of validated events and cell viability EC_50_ values indicates that splicing changes remain a promising surrogate for PRMT5 activity. Moreover, we confirmed these events in an in vivo *MTAP*-deleted GBM CDX model, highlighting their translational potential.

Currently, SDMA levels are typically assessed in serum or via IHC on fixed tissue (41–47). Although informative, IHC-based assays are semiquantitative, often constrained by limited dynamic range, and global post-translational modifications such as SDMA do not provide the pathway-level resolution necessary for mechanistic interpretation. The development of a complementary, quantitative, and standardized splicing-based biomarker could therefore offer substantial orthogonal value, enhancing the ability to monitor target engagement and guide the clinical development of PRMT5 inhibitors.

In summary, our findings show that PRMT5 inhibition consistently perturbs splicing fidelity in *MTAP*-deleted cells, producing reproducible, lineage-agnostic events that are dependent on PRMT5 activity. These events may serve as a robust pharmacodynamic signature, complementing SDMA, and could ultimately help catalog PRMT5 inhibitor efficacy both in vitro and in vivo.

## DATA AVAILABILITY AUTHOR CONTRIBUTIONS

Conceptualization, M.R.T, and K.J.B.; Methodology, M.R.T., S.R.M, and S.L.; Formal Analysis, M.R.T., and S.R.M.; Data Acquisition, M.R.T.; Writing – Original Draft, M.R.T.; Writing – Review and Editing, M.R.T., L.C., and K.J.B.; Visualization, M.R.T; Supervision, L.C., and K.J.B.

## ACKNOWLEDGEMENTS

We thank Dr. Serge Gueroussov for splicing analysis assistance.

## List of Figures

**Supplemental Figure 1 – TNG908 pharmacodynamic effects are histology agnostic**

**A:** SDMA immunoblots following a 3-day treatment with TNG908 at cell line-specific EC_20_ and EC_50_ concentrations from (Figure 1A).

**Supplemental Figure 2 – Endogenous alternative splicing does not predict TNG908 sensitivity**

**A:** Correlation of CCLE reported total ASE’s from *MTAP*-deleted to the cell line-specific EC_50_.

**B:** Correlation of CCLE reported ASE’s for SE, MXE, A3SS, A5SS and RI from *MTAP*-deleted to the cell line-specific EC_50_.

**Supplemental Figure 3 – PRMT5 inhibition induces alterations in alternative splicing**

**A:** TNG908 viability assessment as measured in a 7-day assay using LN-18 (GBM) *MTAP*-isogenic cell lines. Lines were created by reintroducing exogenous MTAP into an *MTAP*-deleted cell line. Data are presented as mean ± SD.

**B:** TNG908 viability assessment as measured in a 7-day assay using either Miapaca2 (PDAC), HCT116 (CRC), or A549 (NSCLC) *MTAP*-deleted lines. Data are presented as mean ± SD.

**Supplemental Figure 4 – Subset of validated events are specific for PRMT5 inhibition.**

**A:** Determination of maximum effect (Amax) in 34 cancer cell lines representing multiple cancer lineages including melanoma, pancreatic adenocarcinoma, mesothelioma, NSCLC, cholangiocarcinoma, and glioblastoma following 7-days TNG908 treatment. Cell lines are colored by MTAP status.

**B:** Normalized SDMA levels following a 1-day exogenous administration of MTA in HAP1 *MTAP*-WT cells.

