## Supplementary figures and images for "Identification of a lineage-agnostic splicing signature caused by PRMT5 inhibition"

### Supplemental Figure 1

#### Supplemental - 1

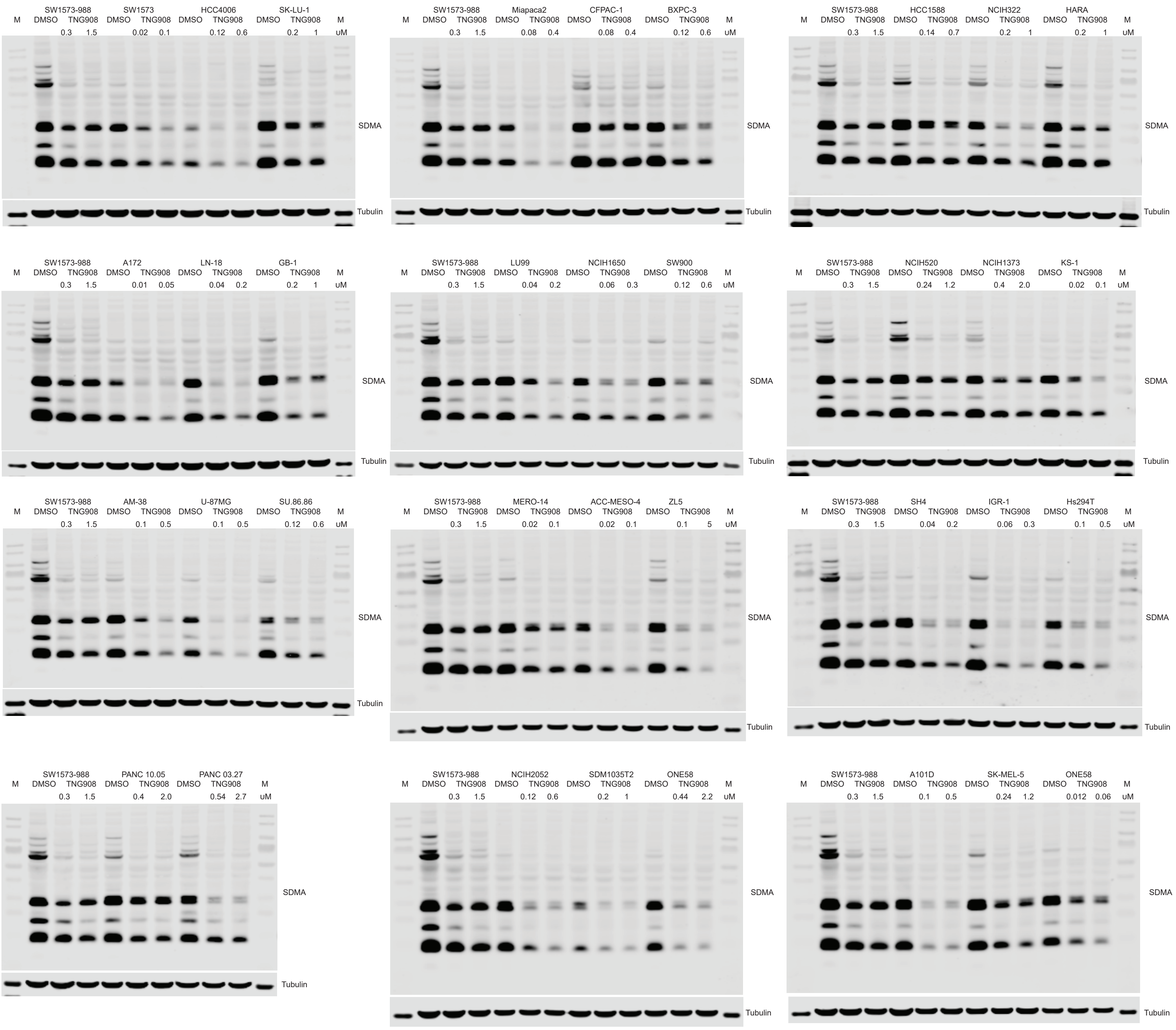

### Supplemental Figure 2

#### Supplemental - 2

## A

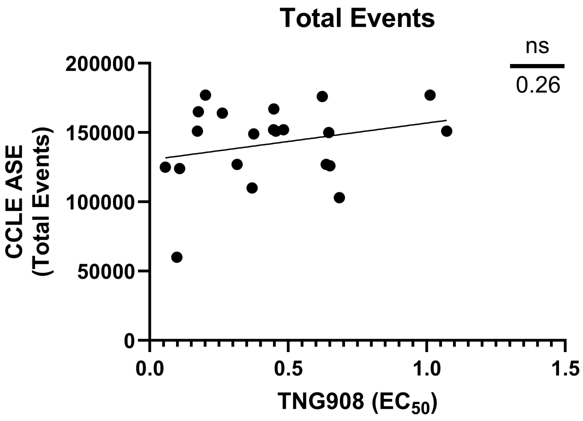


## B

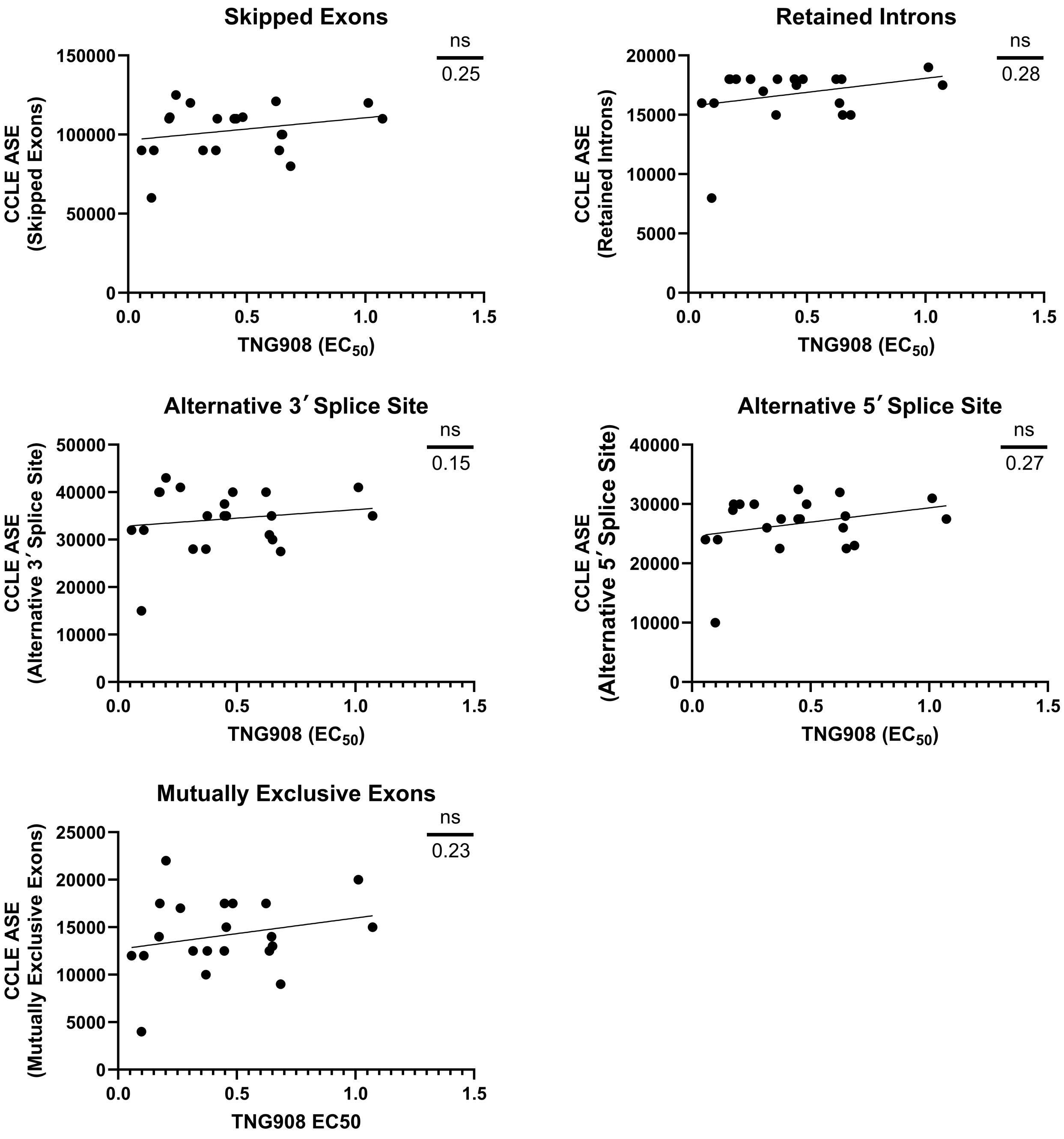

### Supplemental Figure 3

##### Supplemental - 3

**A**

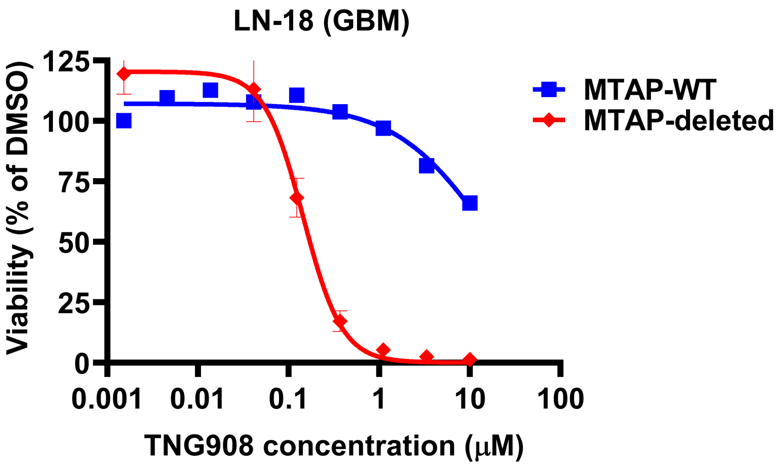


**B**

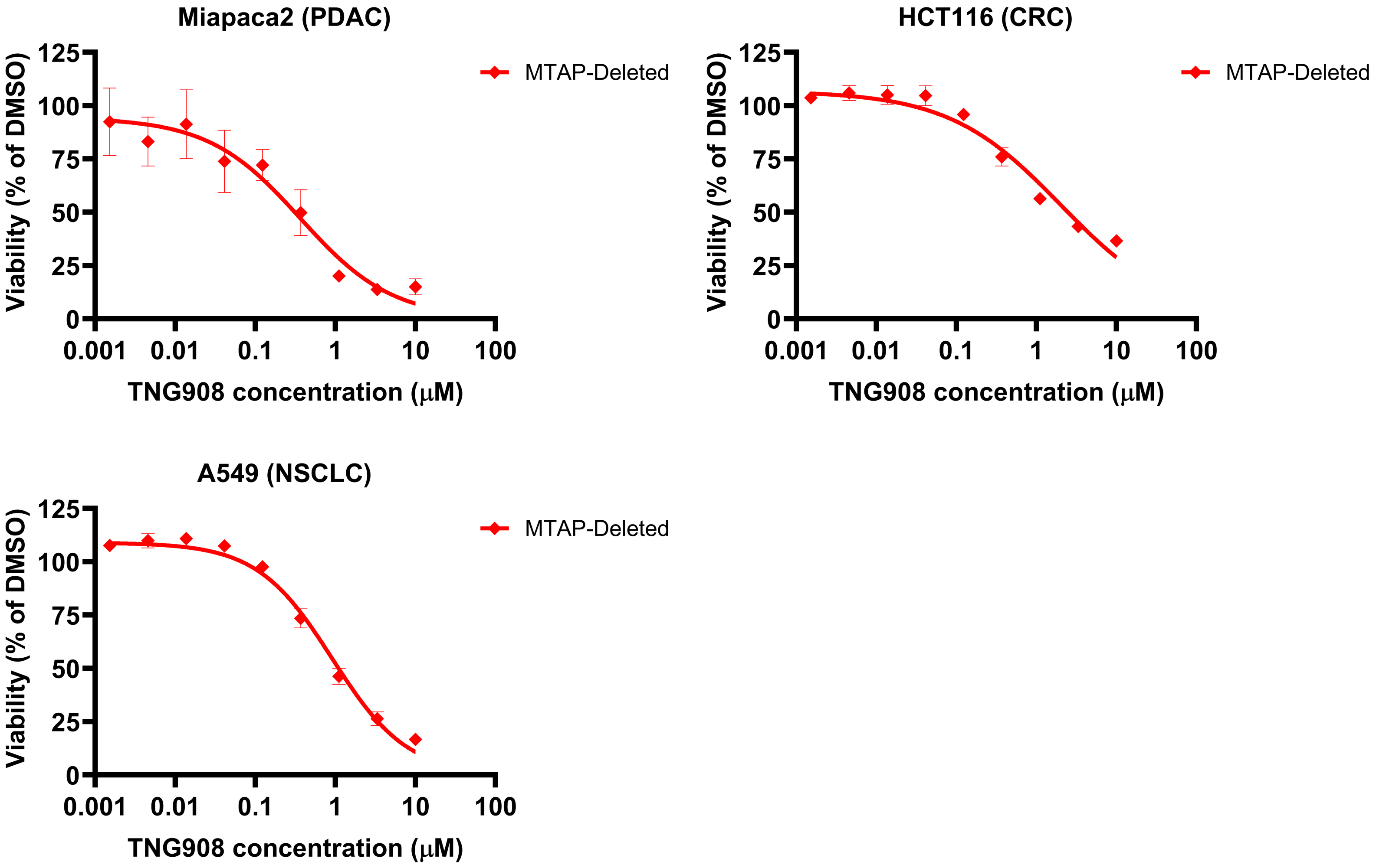

### Supplemental Figure 4

##### Supplemental - 4

**A**

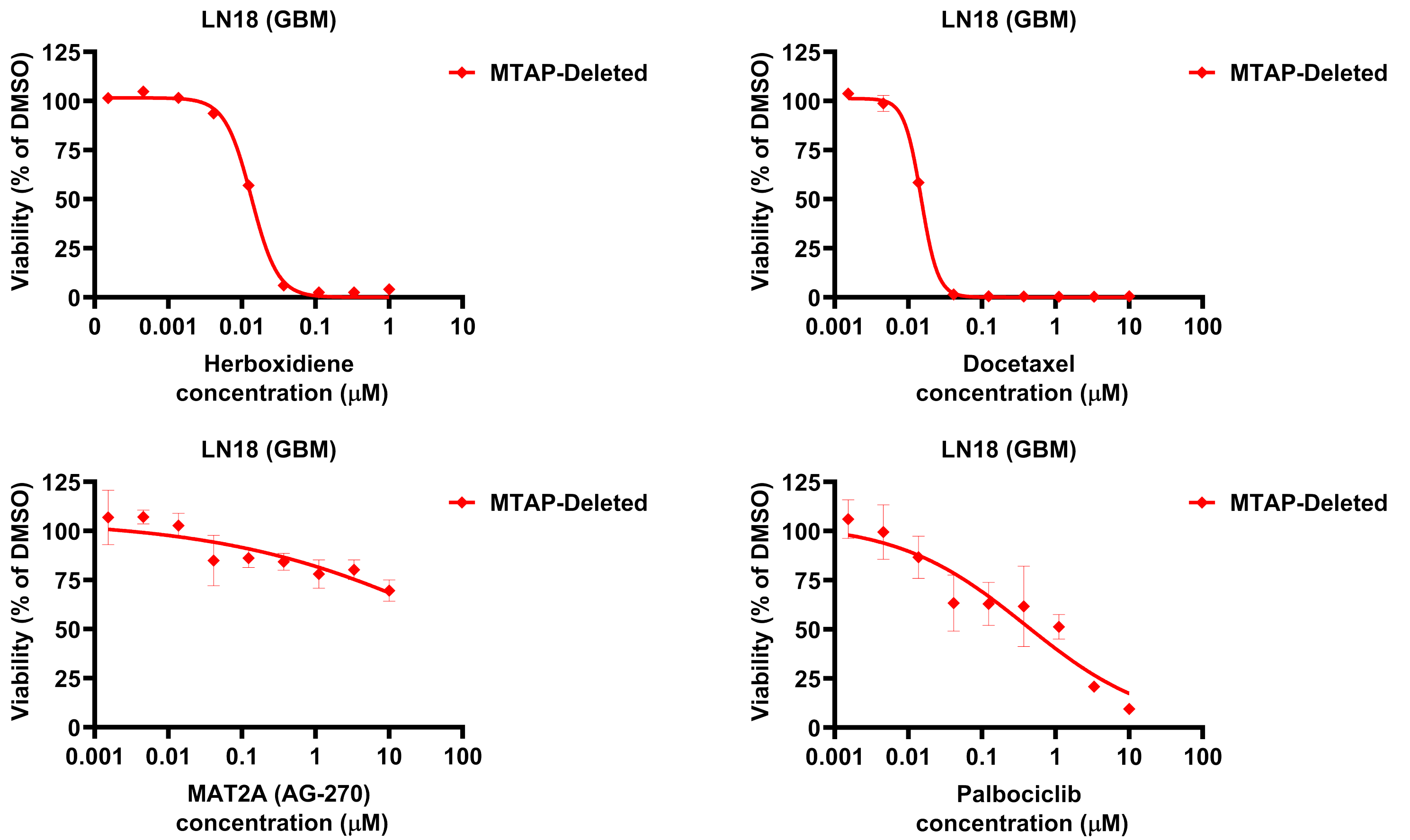


**B**

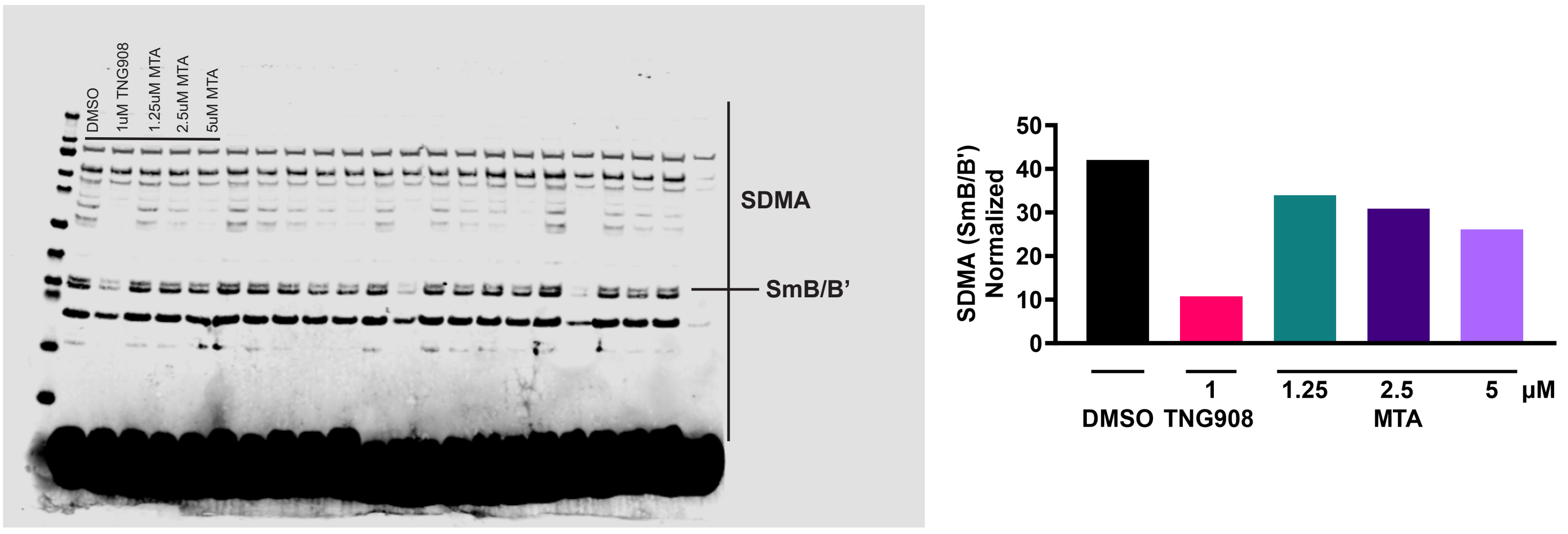
